# Insights into the protective mechanisms of N-acetyl cysteine against D-Galactosamine induced acute liver injury in rabbits

**DOI:** 10.1101/2025.08.24.672039

**Authors:** Neeta Karont, Taru Verma, Adhira Kolangara, Adhya Lakshmi, Mahesh Gopasetty, Jagadeesh Gopalan, Akshay Datey

**Affiliations:** Ykrita Life Sciences Pvt. Ltd. Bangalore, INDIA; Department of Aerospace Engineering & Centre for Excellence in Hypersonics, Indian Institute of Science, Bangalore, INDIA

**Keywords:** Acute liver failure, acute liver injury, D-galactosamine, survival, LFTs, PT-INR, coagulopathy, telemetry

## Abstract

Pre-clinical studies in test animals like rodents, rabbits and pigs are extremely important in establishing the efficacy of therapeutic interventions ranging from formulations to medical devices. Diseases like acute liver failure (ALF) are although rare, but most often culminate into a situation where the patient needs an emergency liver transplant. Currently, not many medications are available to manage such patients. Out of the very few available options, N-acetyl cysteine (NAC) is the clinician’s drug-of-choice. Administration of NAC has shown improvements in patient’s survival in the hospitals. Yet the mechanism and its systematic investigation in pre-clinical models like rabbits remains undone. In this report, we establish a drug-induced, controlled-ALF in rabbits using D-galactosamine (D-Gal). We also test the efficacy of NAC in this model system. We support the *in-vivo* findings with detailed *in-vitro* assays where we attempt to understand the mechanisms of the prophylactic usage of NAC against D-Gal induced ALF.

## Introduction

Acute Liver Failure (ALF) is a rare and fatal condition, characterized by the sudden loss of hepatic function within 7 to 28 days due to hepatocyte cell death in previously healthy individuals ^1^. The incidence of ALF in developed countries is 6 cases per million population per year ^2,3^. Similar data in the Indian context largely remains unreported due to lack of data recording infrastructure. Extrapolating the data from developed nations to the current Indian population, 8496 cases of ALF are estimated in 2023. However, this number is an underestimate, and the incidence is expected to be much higher ^4,5^. According to King’s College Criteria, ALF is defined as a clinical syndrome where encephalopathy occurs at an interval of 8 to 28 days after the onset of jaundice ^1^. ALF is diagnosed with an array of liver function tests (LFTs). Precisely, prothrombin time (PT) > 100 seconds, INR (International Normalized Ratio) ≥1.5, serum creatinine 300 μmol/L, elevated alanine transaminase (ALT), aspartate transaminase (AST) are the biomarkers ^6^. The most common etiology of ALF is acetaminophen overdose in the United States, followed by non-acetaminophen drug induced liver injury (NAI-ALF). Viral hepatitis contributes the most to the non-drug induced cases of ALF in developing nations like India ^7^.

Currently, there are no established treatments for NAI-ALF apart from orthotopic liver transplantation (OLT) ^8^. Despite the high cost and limited donor availability, the success rate of OLT is 70% over 10 years ^9^. Failure is attributed to conditions like graft rejection, graft failure or post-surgery complications like infection. Therefore, it is important to develop strategies which are minimally invasive and cost-effective in treating ALF. The current non-surgical treatment for ALF includes usage of hepatoprotective drugs like N-acetyl cysteine (NAC), glutathione, glycyrrhizin acid preparation, bicyclol, carnitine, polyene phosphatidylcholine, etc. ^10^. These drugs improve liver function, promote liver cell regeneration and/or enhance detoxification. Among these, NAC is the most promising as it has shown improvement in LFP, shorter hospital stays and increased survival in clinical settings ^10^. NAC, a derivate of a sulphur-containing amino acid, L-cysteine, is a well-tolerated mucolytic drug ^11.^ Once ingested, NAC is deacetylated in the liver and acts as a potent antioxidant. It stimulates glutathione synthesis and thereby facilitates detoxification ^11^. Owing to its excellent free radical scavenging capacity, NAC is under extensive investigation to treat conditions associated with oxidative stress ^11^. One such implication is the use of NAC in treating acetaminophen induced in patients and laboratory animal models ^12^. Its role in non-acetaminophen induced ALF (NAI-ALF) is elusive ^13^. Although the efficacy and mechanism of NAC is unclear, it is routinely used as non-standard care therapy to manage ALF arising out of multiple etiologies ^14^.

In this study, we have evaluated the protective action of NAC in managing drug-induced ALF. To test this hypothesis, we employed the D-Galactosamine (D-Gal) induced ALF model. D-Gal, a derivative of galactose, is a potent hepatotoxic agent that causes cell death by necrosis and apoptosis ^15^. High levels of galactokinase and galactose-1-uridyltransferase facilitate tissue specific uptake of D-Gal in liver ^16^. D-Gal depletes the pool of uracil nucleotides, thereby inhibiting RNA synthesis ^17^. In addition, D-Gal also contributes to excessive oxidative stress leading to cell death ^18^. Therefore, D-Gal is commonly used to experimentally induce ALF in various animal models. Pigs are the most accepted models to study liver diseases due to their anatomical and physiological similarities to humans ^19^. Rodent models of ALF are common due to their small size, low cost and availability. However, the main criterion of ALF assessment is restricted to the elevated transaminases in rodents. Coagulopathy and kinetics of ALF progression in rodents cannot be monitored due to limited blood volume ^20^. Rabbits, on the other hand, have an intermediate size that allows ready blood sampling, and show an intermediate sensitivity to D-Gal^21^. Their immune system also resembles humans more than rodents ^22^. Therefore, the rabbit model is better suited for studying the kinetics of D-Gal induced liver injury and protective effects of NAC that remains unreported in rabbits till date.

## Materials & Methods

### Chemicals

D-Galactosamine-Hydrochloride (SRL-14725) was used for the induction of acute liver failure (ALF) and to study the *in-vitro* toxicity. N-Acetyl L-cysteine (A9165) and DCFH-DA (287810) were purchased from Sigma-Aldrich. ATP determination kit (A22066) was obtained from Thermo-Scientific and liquiplastin reagent for prothrombin time (PT) was purchased from Tulip Diagnostics.

### Animals

New Zealand White Rabbits were acquired at the age of 2 months and were given an acclimation period of 7 days before the initiation of any procedure. Animals were housed at Radiant Research Centre, an ISO certified Contract Research Organization (CRO) accredited with NABL, India. All the animals had free access to food and water and were maintained in controlled temperature (21°C-25°C) rooms with 12 h light and dark cycle. During the quarantine period the animals were treated with 0.4 mg/kg ivermectin (subcutaneous injection) and 10 mg/kg enrofloxacin (IM injection) every alternate day for 7 days. Animals weighing 1.5 kg-2.5 kg and with basal blood biochemistry in the range described in Table 1. were included in the experiment (Figure 1A).

**Figure 1:**
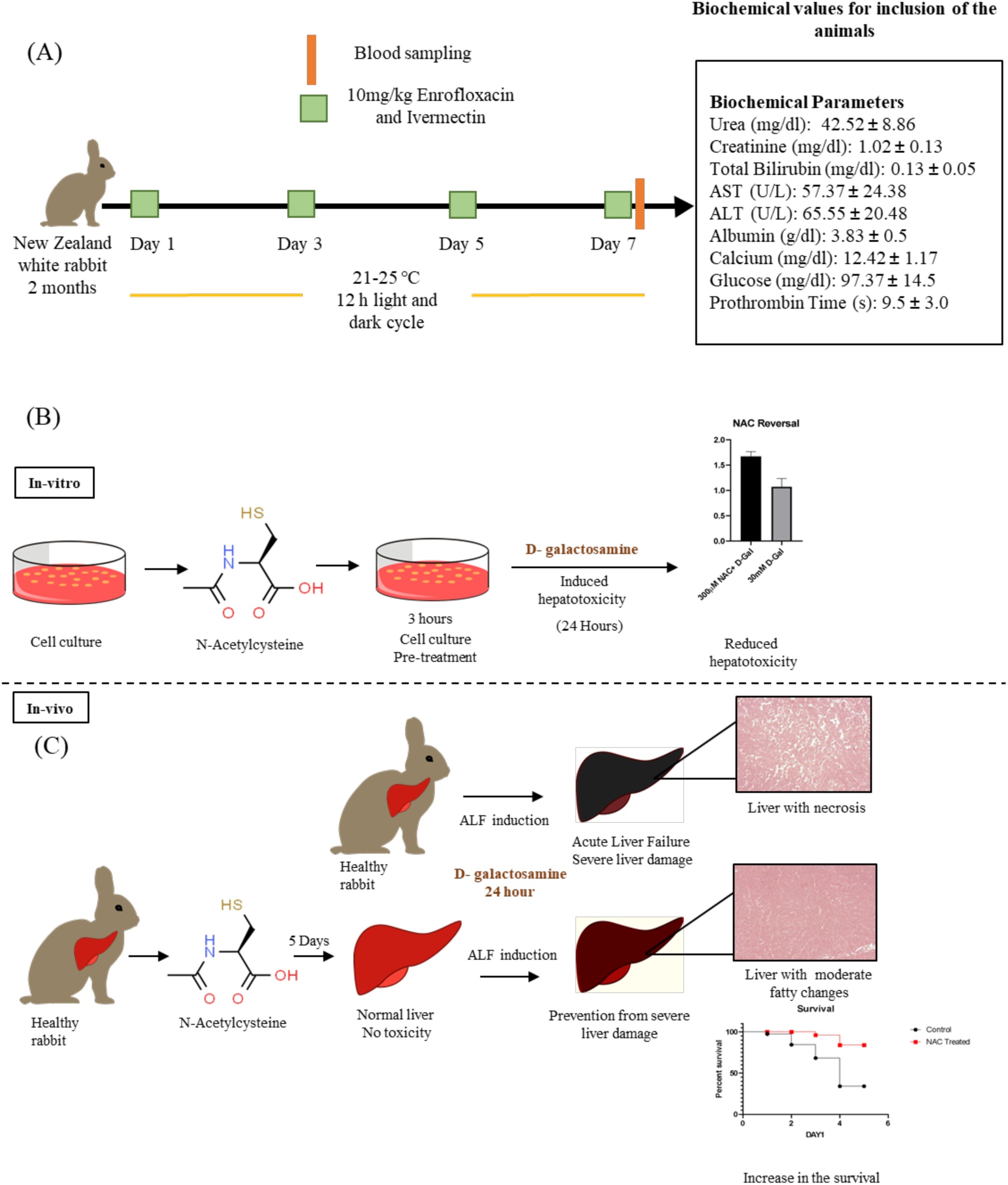
Animal model selection and experimental strategy. (a) Animal quarantine protocol and inclusion criteria for the experiment. (b) Experimental strategy for exploring the protective effects of NAC in-vitro and (c) in-vivo in rabbits.

**Table 1:** Normal range of biochemical parameters in New Zealand White rabbits.

| Biochemical Parameter | Value |
| --- | --- |
| Urea (mg/dl) | 42.52 ± 8.86 |
| Creatinine (mg/dl) | 1.02 ± 0.13 |
| Total Bilirubin (mg/dl) | 0.13 ± 0.05 |
| AST (U/L) | 57.37 ± 24.38 |
| ALT (U/L) | 65.55 ± 20.48 |
| Albumin (g/dl) | 3.83 ± 0.5 |
| Calcium (mg/dl) | 12.42 ± 1.17 |
| Glucose (mg/dl) | 97.37 ± 14.5 |
| Prothrombin Time (s) | 9.5 ± 3.0 |

### Cell culture

Hepatocyte cell line HepG2(ATCC HB-8065) was used as a model for all *in-vitro* experiments. HepG2 cells were cultured in DMEM high glucose (Sigma Aldrich D6429) supplemented with 10% heat inactivated FBS (Gibco 10270-106) and 1% penicillin-streptomycin mix (Sigma Aldrich P4333).

### Blood Biochemistry

Whole blood was collected from the marginal ear vein of rabbits into respective tubes (EDTA, clotting-activated & citrate) and plasma/serum were isolated by centrifugation at 5000 rpm for 10 minutes at room temperature (RT). The samples were analyzed using Agappe Mispa Ace biochemistry analyzer (SGOT, SGPT, bilirubin, urea, creatinine, albumin and calcium). Prothrombin time (PT) was checked using liquiplastin kit from citrate-plasma according to the manufacturer’s instructions.

### NAC treatment *in-vivo*

A total of 6 animals were used in this study, where they were randomly divided into two groups-control and NAC treatment group (n=3). Controls were injected with phosphate buffered saline (PBS) and the NAC group received 500 mg/kg NAC injection IM. Treatment was given every 24 hours for 5 days and the general health of the animals was observed. Blood was sampled every 48 hours for biochemistry analysis (Figure 1C).

### NAC treatment *in-vitro*

For *in-vitro* NAC treatment in HepG2 cell line, 1 x 10^4^ cells/well were seeded in a microtiter plate and were allowed to grow overnight. 50 mM NAC was freshly prepared in PBS and the pH was adjusted to 7.4. 100µl of NAC containing medium was added to the cells at the concentrations ranging from 100mM - 500mM. Cells were then incubated at 37°C for 24 hours (Figure 1B).

### Establishing D-Galactosamine induced ALF *in-vivo*

Animals were randomly divided into vehicle-control and D-Galactosamine-treatment groups (n=3). Basal blood biochemistry was performed on all animals before any treatment. Different concentrations of D-Galactosamine were prepared in 5% dextrose and pH was adjusted to 6.8-7. Rabbits in the treatment group received dose concentrations of 100mg/kg, 200mg/kg, 300mg/kg, 400mg/kg, 450mg/kg, and 500mg/kg IV, while vehicle controls received 5% dextrose solution. Post injections, animals were monitored for the onset of ALF by routine blood sampling.

### Estimating D-Galactosamine LD_50_ in-vitro

For determination of LD_50_ in HepG2 cell line, 1×10^4^ cells were seeded in microtiter plate and were allowed to grow overnight. D-Galactosamine was prepared in PBS and adjusted to pH 7.0 - 7.4. 100µl of D-Gal containing media was added to the cells at the dose concentrations of 10 – 100 mM. The cells were incubated for 24 hours, and toxicity was determined using phase contrast imaging and MTT assay.

### Induction of Acute Liver Failure (ALF)

To induce ALF, animals were injected with a single IV dose of 500 mg/kg D-Galactosamine, after 5 days of NAC or PBS pre-treatment. Blood was analyzed every 24 hours for 7 days to monitor the progression of ALF. This continued till the animals recovered or succumbed to ALF.

### Induction of Hepatotoxicity *in-vitro*

D-Galactosamine was prepared as described above. All the cells were treated with 20 – 30 mM D-Gal for 24 hours. Cell viability was assayed using phase contrast imaging and MTT assay.

### Necropsy and Histology

Animals were euthanized using ethically approved IV injection of pentobarbital. During necropsy, liver was harvested and fixed in 4% paraformaldehyde. The organ was dehydrated and embedded in the paraffin block. 5µm thick tissue sections were taken using a microtome and the sections were processed for hematoxylin and eosin staining (H&E). Slides were then observed under a bright field microscope and the images were analyzed using ImageJ software. Histopathological scoring was performed by an external pathologist. The histological examination of liver was performed on animals from the vehicle control group, when they became moribund, while animals in NAC treatment group were euthanised as per the ethical guidelines, after they survived the expected period of 30 days.

### MTT Assay

Cell viability was estimated using MTT assay. Briefly 5 x 10^4^ cells/well were seeded in a 24 well plate and incubated overnight. D-Gal and/or NAC treatment was done as described earlier. The cells were washed twice with PBS. MTT (3-(4,5-dimethylthiazol-2-yl)-2,5-diphenyl tetrazolium bromide) was added at a final concentration of 0.5mg/ml after the treatment and the cells were incubated in MTT medium for 4 hours at 37°C in dark. Formazan crystals were solubilized using dimethyl sulfoxide (DMSO). Absorbance was read at 570 nm with DMSO as reference.

### ROS estimation

Quantification of reactive oxygen species (ROS) was done using 2’,7’ dichlorofluorescin diacetate (DCFH-DA) staining. Briefly, 5×10^4^ cells per well were seeded in a 24-well plate and allowed to grow overnight. 20mM D-Gal treatment was done for 3 hours. Cells were then trypsinized and cell number was estimated using a hemocytometer. Cell suspension was stained using 10mM DCFH-DA for 30 minutes at 37°C in dark. Cells were washed (2x) using PBS by centrifugation at 2000 rpm for 5 minutes at RT. Cell pellet was resuspended in PBS and the fluorescence was read at λ_ex_ 485 nm and λ_em_ 530 nm using Tecan Infinite PRO 200. The fluorescence was normalized by cell count and was expressed as relative fluorescence unit (RFU).

### ATP Assay

Intracellular ATP was assayed using ATP Determination Kit (Invitrogen). 5×10^4^ cells per well were cultured in 24-well plate and allowed to grow overnight. Cells were treated with 20 mM D-Gal for 24 hours as described above. Cells were then trypsinized and were enumerated using a hemocytometer. Cells were washed by centrifugation at 2000 rpm for 5 minutes (2x) using PBS and the pellet was suspended in 100µl deionized water. Boiling lysis was performed using a thermal cycler set at 99°C for 10 minutes. Cell lysate was immediately plunged into ice. Lysate was centrifuged at RT at 1000 g for 1 minute and 10µl of the supernatant was assayed using the kit. Luminescence readings were recorded for all samples using Tecan Infinite PRO 200. The total ATP concentration was determined using the ATP standard curve and expressed as ATP per cell.

### Real time monitoring of vital parameters

Animal vital parameters were monitored using wired telemetry system PowerLab C from AD Instruments. Data was acquired using LabChart 8 software. For temperature measurement a rectal probe was used. ECG was acquired by placing the bio-amp probes subcutaneously in the animal forelimbs and hind-limbs. Heart rate was derived from ECG and respiration rate was obtained by placing the bridge amp pressure probe under the diaphragm of the animal. For analyzing the ECG data, the cyclic measurements were converted into a single event after setting the R-R interval to 100 milliseconds. All the events acquired throughout the measurement were averaged and were represented as a single event.

### Statistics

Data analysis and graphical representation was done using GraphPad Prism (Version 9). Survival between two independent groups was analysed using Log-rank test while the independent groups were analysed using unpaired t-test. Significance is represented as p*<0.05, p**<0.01, p***<0.005. ‘N’ indicates the number of independent experiments and ‘n’ represents the number of replicates.

## Results

### Evaluation of D-Galactosamine induced toxicity and protective effects of NAC in a hepatocyte cell line

D-Galactosamine, a galactose derivative, is a hepatotoxic agent that causes hepatocyte cell death by necrosis and apoptosis. To understand the hepatotoxic effect of D-Gal, we performed a dose dependent study on HepG2 cell line *in-vitro*. Cells were treated with D-Gal for 24-hours and we observed a dose dependent lethality as shown in the phase contrast image panel (Figure 2A). The dose response curve was generated using MTT assay and the LD_50_ value was estimated to be around 23.89 mM for HepG2 cell line (Figure 2B). Therefore, the concentration from 20-30 mM was used for all the subsequent experiments *in-vitro*. To understand the mechanism of cytotoxicity, we measured the levels of intracellular ATP and reactive oxygen species in the cells. We observed that the cell number and total ATP in D-Gal treated samples had reduced, while there was no significant change in ATP per cell. Therefore, D-Gal does not perturb ATP pool in the cytoplasm. Hence, cytotoxicity may not be attributed to any changes in the ATP pool (Figure 2C). Regardless of the etiology, oxidative stress and inflammation are crucial pathogenetic markers in liver diseases ^23^. D-Gal has been reported to cause hepatotoxicity by increasing the reactive oxygen species. We also found that D-Gal treated samples had a 2-fold increase in intracellular ROS after 3 hours of exposure, compared to the untreated control group (Figure 2D). Glutathione (GSH) is the most abundant thiol containing antioxidant in the cell, which is synthesised in the presence of cysteine ^11^. During ALF, there is a depletion in levels of GSH, which causes an increase in oxidative stress levels ^24^. Currently NAC is being studied in diseases associated with increased oxidative stress or decreased GSH levels ^11^. NAC acts as a precursor of GSH, it can be passively transported across the cell membrane and is deacylated to form cysteine in the cytoplasm ^25^. The efficacy and mechanism of NAC in inhibiting cytotoxicity in hepatocytes is unreported. To address this, a dose dependent study was conducted on HepG2 cell lines *in-vitro*. NAC alone did not affect the viability of hepatocytes. Normal cell morphology was observed post 24 hours of NAC treatment as shown in the phase contrast image panel (Figure 2E). Cell viability was assessed to check the adverse effect of NAC using MTT assay and was found to be insignificant (Figure 2G). Previous reports have shown that NAC pre-treatment can protect against various hepatotoxic compounds like DDT and its metabolites ^26^, but the dose dependent efficacy remains unexplored. To assess the prophylactic effects of different doses of NAC against D-Gal, the cells were treated with NAC for 3 hours. Various doses of NAC were used to determine if these effects were dose dependent as shown in the phase contrast image panel Figure 2F. NAC pre-treatment protected the cells against D-Gal toxicity and the protective effect showed saturation at 100µM (Figure 2H). This indicates that the efficacy of protection did not increase with higher doses of NAC. Overall, these results indicate that NAC has hepatoprotective effect *in-vitro,* with a minimum dose of 100µM being sufficient for inhibition of drug induced toxicity.

**Figure 2:**
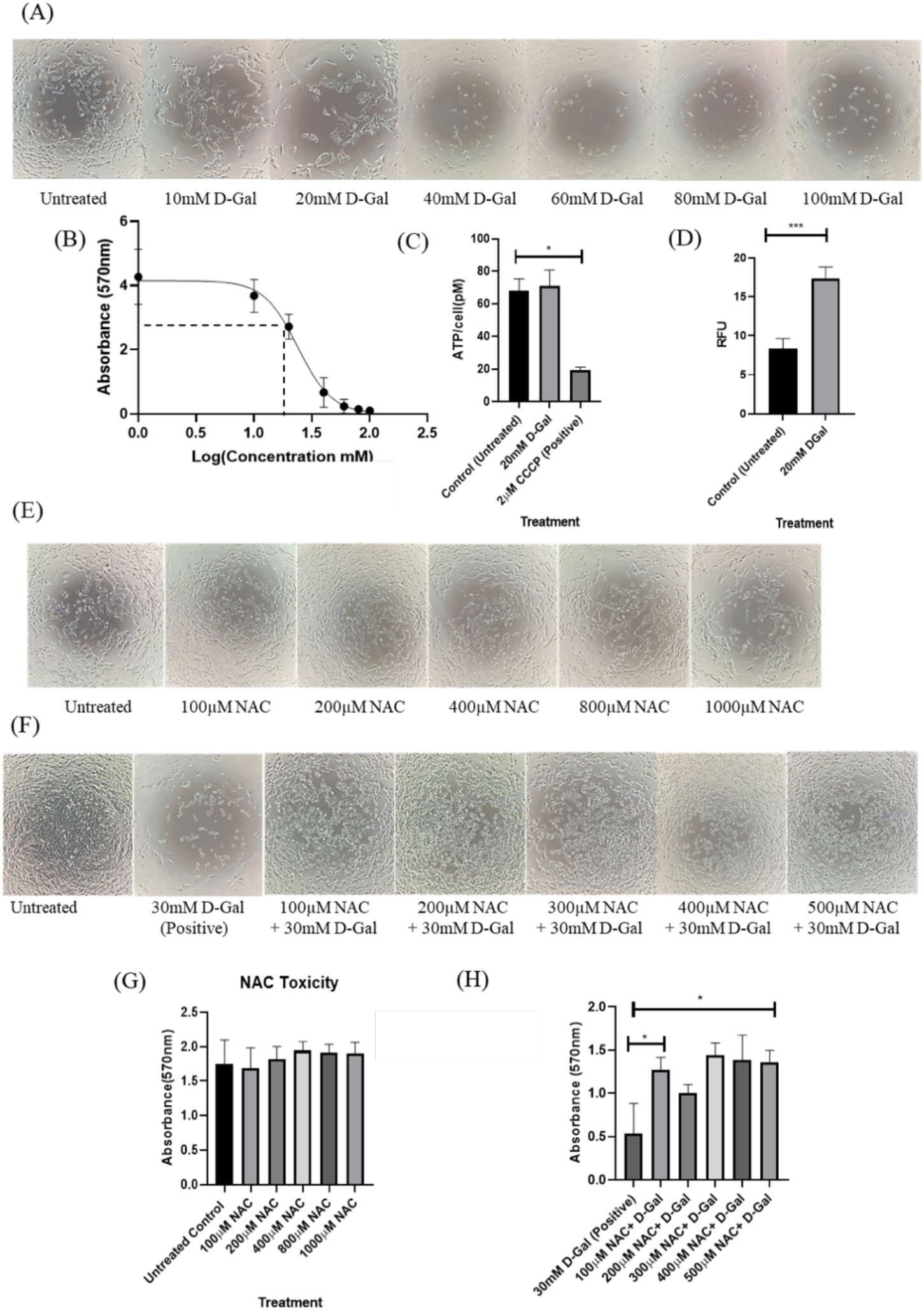
Evaluation of D-Galactosamine (D-Gal) toxicity and protective effects of NAC in hepatocyte cell line. (A) Representative phase contrast images of HepG2 cells after 24 hours of D-Gal (10-100 mM) treatment. Linear relationship between the dose and cell viability was observed. (B) Dose response curve of HepG2 cells treated with D-Gal (10-100mM) for 24 hours determined using MTT assay (N=3, n=3). (C) Estimation of ATP levels in HepG2 cells treated with D-Gal (LD_50_) for 24 hours. The total ATP was estimated using ATP determination kit and the chemiluminescence was recorded using plate reader. Cells treated with 2 µM CCCP for 20 minutes were used as a positive control (N=2, n=3). (D) Graphical representation of reactive oxygen species in HepG2 cells treated with D-Gal (LD_50_) for 3 hours determined using DCFH-DA staining (N=3, n=3). (E) Represents the phase contrast images of the HepG2 cells treated with NAC (100-1000µM) for 24 hours. (G) Depicts the cell viability (MTT assay) after 24 hours of NAC treatment (100-500 µM) in HepG2 cells (N=3, n=3). (F) Represents the phase contrast images of HepG2 cells that received a pre-treatment with 100µM-500µM NAC for 3 hours followed by D-Gal treatment for 24 hours. (H) Graphical representation of cell viability determined by MTT assay in HepG2 cells treated with NAC followed by D-Gal as described above in figure legend F (N=1, n=3). All the graphs were analysed using 2 tailed unpaired t-test; p*<0.05, p**<0.01, p***<0.005.

### Establishment of D-Galactosamine Induced Controlled Acute Liver Failure

D-Galactosamine is commonly used for screening hepatoprotective drugs *in-vivo*. Among various animal models, the sensitivity to D-Gal is not linear. Smaller animals like rats have low sensitivity (1.5-2g/kg) ^27^whereas large animals like pigs have been reported to have high sensitivity (0.2-0.4 g/kg) ^28^. Rabbits are moderately sensitive to D-Gal and their appropriate body weight allows frequent blood sampling as compared to rats/mice ^21^. Therefore, establishing a controlled liver injury model in rabbits might prove beneficial in testing therapeutic interventions. Here we sought to establish a closely titrated dose-dependent lethal model of D-Gal induced ALF in rabbits. A single dose of D-Gal at different concentrations (100-500 mg/kg) was administered IV, and the onset of ALF was monitored by analysing LFTs (Figure 3A & 3B). We observed a rapid increase in the levels of aminotransferases in a dose-dependent manner. Animals injected with 450 and 500 mg/kg D-Gal showed a marked increase in ALT and AST after 24 hours of D-Gal injection (Figure 3D and 3E). D-Gal treatment caused anorexia, lethargy and fur loss which are indicative of ALF. In addition, elevations in INR and bilirubin and reduction of serum albumin (Figure 3F, 3G and 3H) also confirmed the severity of ALF.

**Figure 3:**
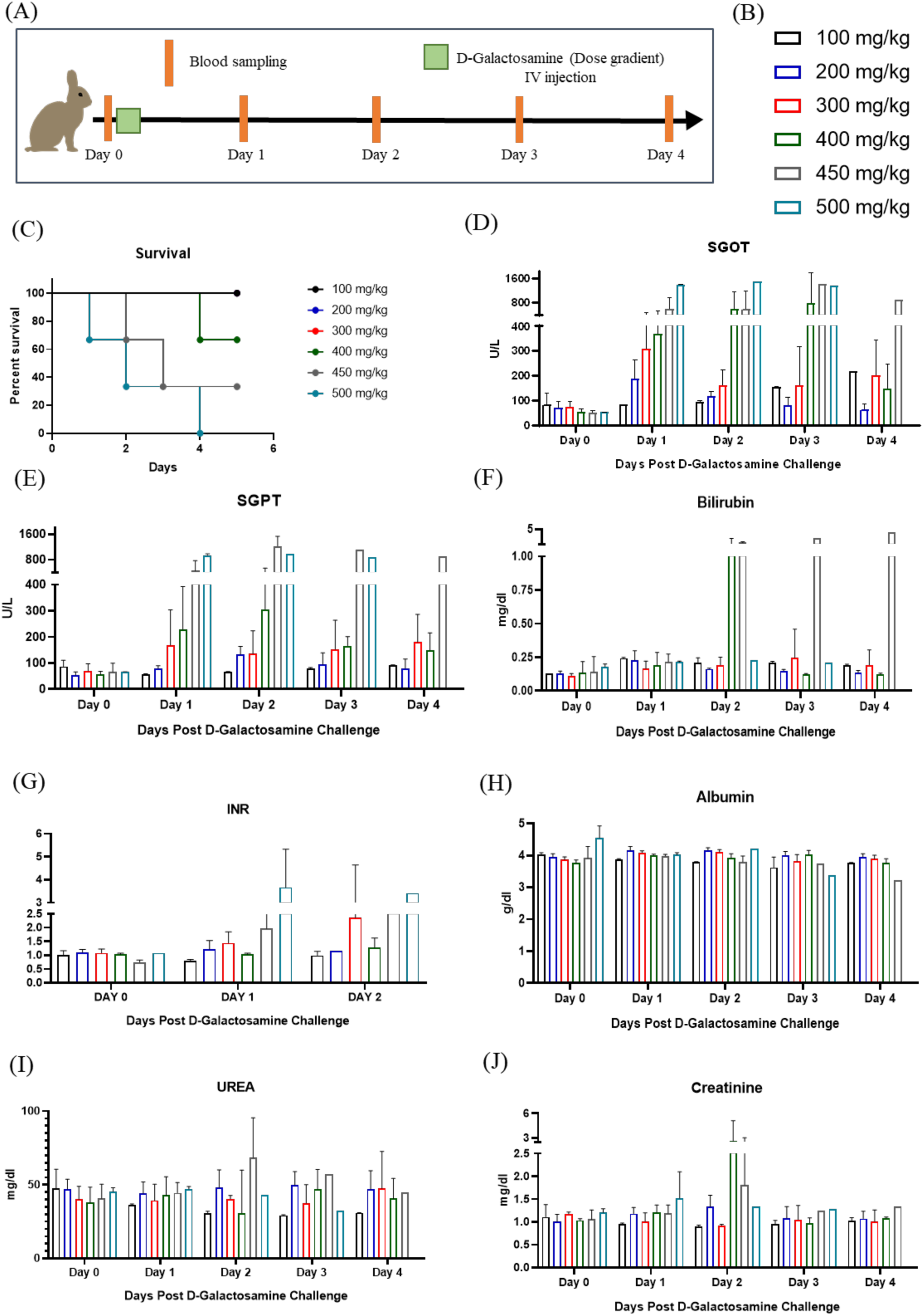
Establishment of D-Galactosamine induced controlled liver injury in rabbits. (A) Schematic representation of experimental strategy for establishment of controlled ALF in rabbits. (B) Colour scheme depicting the various doses of D-Gal (C) Graphical representation of survival after the rabbits were injected with different doses of D-Gal IV (100-500 mg/kg). (D), (E), (F), (G) and (H) Represent the liver function markers namely SGOT, SGPT, Bilirubin, INR and Albumin every 24 hours after injection of various doses of D-Gal. (I) & (J) Represent the kidney function markers namely urea and creatinine every 24 hours after D-Gal injection. All the figures represent pooled data from N=1; n=3.

We observed a linear relation between D-Gal dose and mortality. The dose groups from 100-300 mg/kg did not show any mortality and were able to recover from liver damage. At 450 and 500 mg/kg D-Gal, the animals showed high mortality with mean survival of 33.3 % and 0% respectively (Figure. 3C). D-Gal dose of 500 mg/kg proved lethal as all the animals succumbed to ALF within 4 days post injection. As shown earlier D-Gal increases free radical generation and oxidative stress. The excess oxidative stress upon D-Gal administration is reported to cause renal dysfunction in rodents ^15^. We also observed that D-Gal had off-target effects on renal function in a dose-dependent manner in rabbits as indicated by high urea and creatinine levels (Figure. 3I and 3J).

### D-Galactosamine causes ALF by inducing hepatocyte cell death via necrosis

Liver injury was characterized by microscopic examination of liver. During necropsy, animals treated with high doses of D-Gal revealed discoloured white patches indicative of necrosis (gross images not shown). The histopathology examination showed cellular damage, necrosis, increased portal and periportal inflammation in D-Gal injected animals compared to their healthy counterpart (Figure. 4A & 4B). This aligns with previous reports that have shown D-Gal to cause hepatic injury with spotty hepatocyte necrosis and inflammation in rodents ^29^. Table 2 and Table 3 show the liver histopathology scoring for 450 mg/kg and 500 mg/kg D-Gal injected animals. Therefore, for further experiments 500 mg/kg D-Gal concentration was used as it proved to be a potent dose for a severe ALF in rabbits.

**Figure 4:**
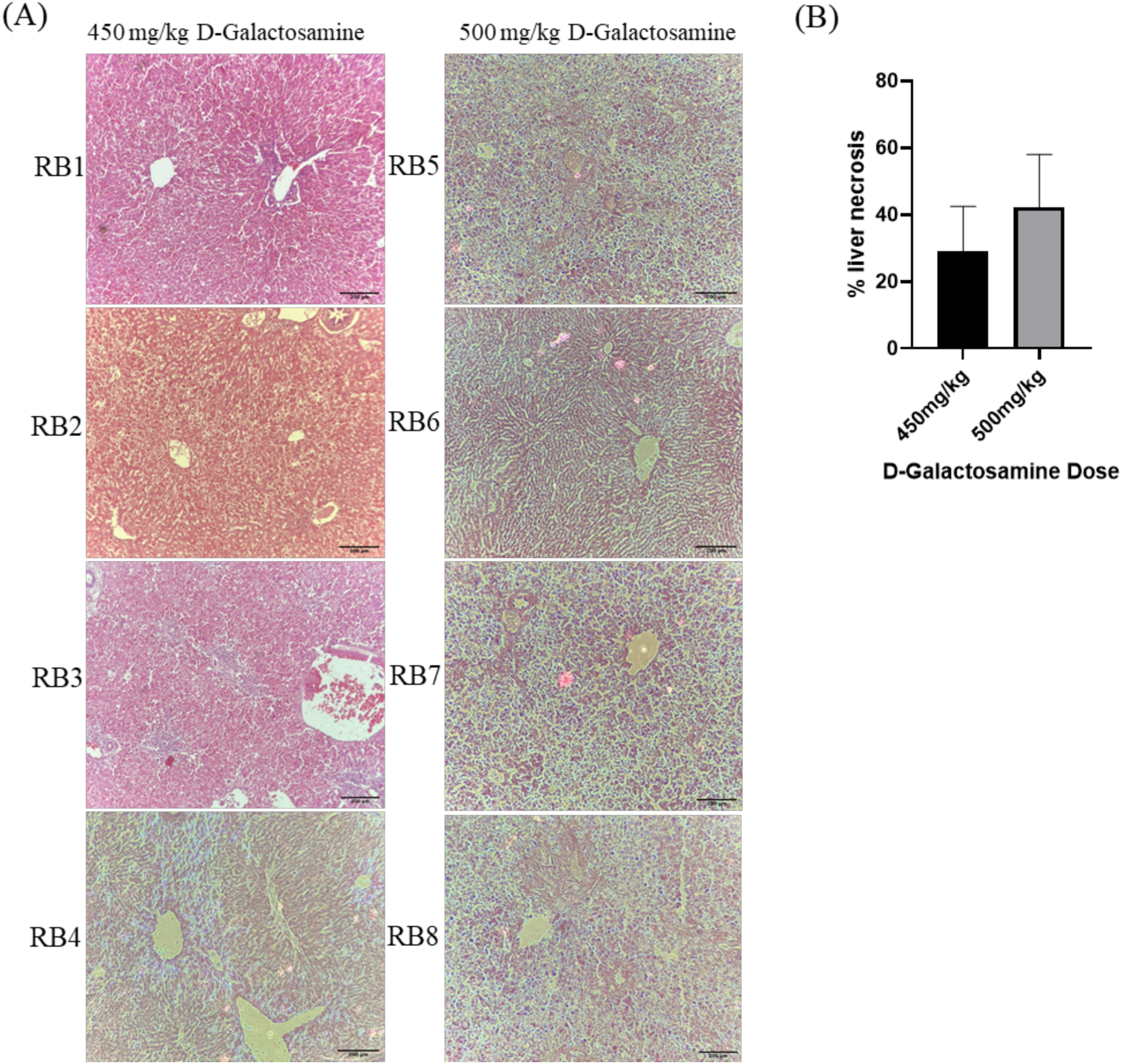
Evaluation of rabbit liver histopathology post D-Galactosamine injection. (A) H&E staining representing hepatocellular structure in 450 mg/kg and 500 mg/kg D-Gal injected animals. The images were captured using a bright field microscope at 10X objective and analysed using Image J. Scale bar = 200 µm (B) Represents the extent of liver necrosis determined by histopathology scoring.

**Table 2:**
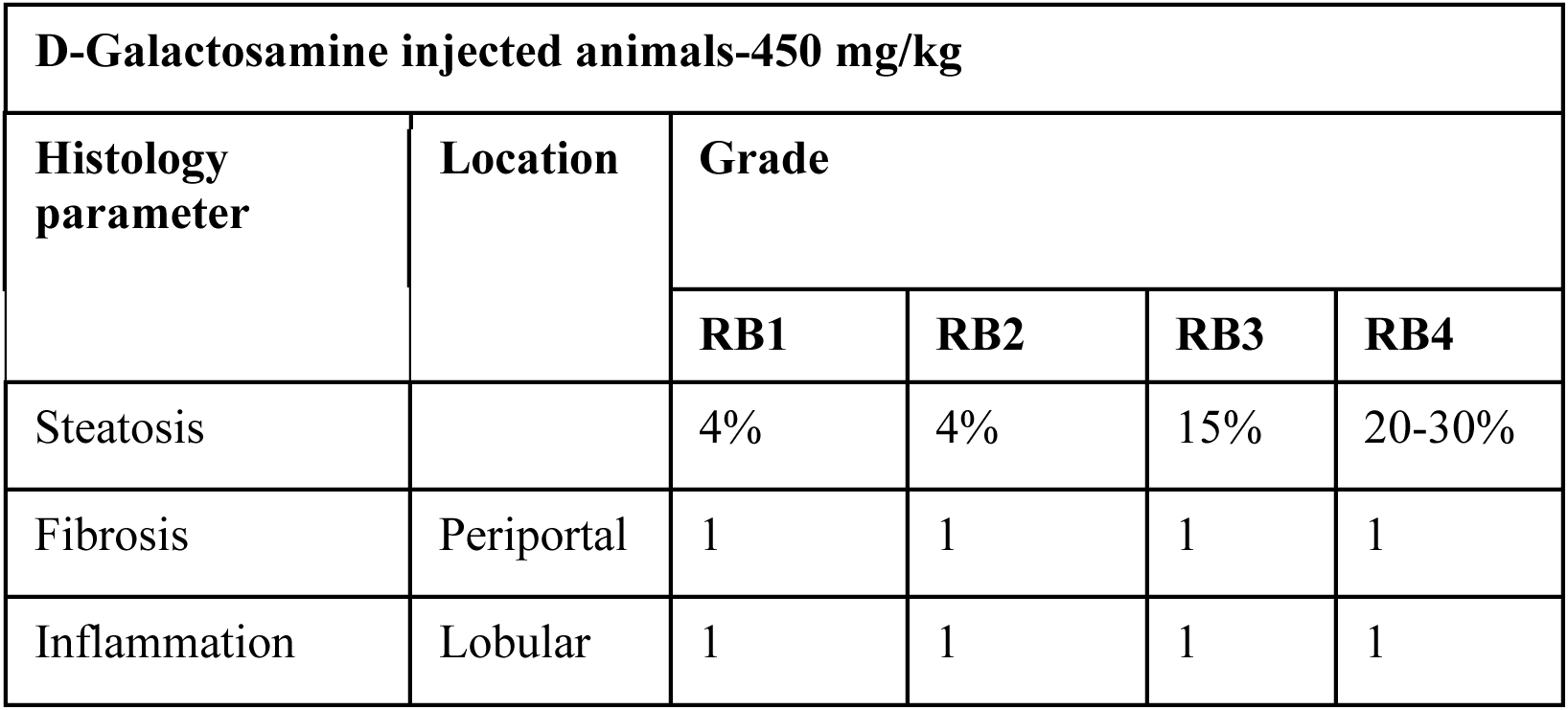
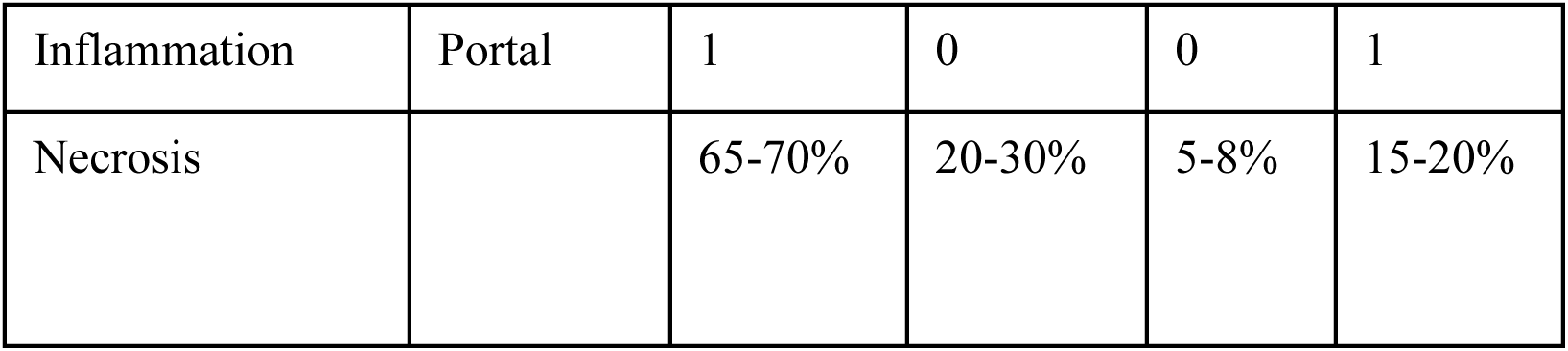
Histopathology Scoring of 450 mg/kg D-Gal injected animals.

**Table 3:** Histopathology Scoring of 500 mg/kg D-Gal injected animals.

| **D-Galactosamine injected animals-500 mg/kg** |  |  |  |  |  |  |  |
| --- | --- | --- | --- | --- | --- | --- | --- |
| **Histology parameter** | **Location** | **Grade** |  |  |  |  |  |
|  |  | **RB5** | **RB6** | **RB7** | **RB8** | **RB9** | **RB10** |
| Steatosis |  | 1% | 1% | 4% | 1%, | 1%, | 4% |
| Fibrosis | Periportal | 1 | 1 | 1 | 1C | 1 | 1 |
| Inflammation | Lobular | 1 | 1 | 1 | 2 | 1 | 1 |
| Inflammation | Portal | 0 | 0 | 1 | 1 | 0 | 1 |
| Necrosis |  | 75-80% | 5-8% | 65-70% | 2% | 55-60% | Absent |

### D-Galactosamine affects animal vital parameters

The knowledge about pathology of ALF in laboratory animal models like rabbits is restricted to LFTs, body weight, hydration, and altered food intake ^30^. To shed light on the impact of ALF on important vital parameters, we monitored previously unknown vitals like body temperature, heart rate, ECG and respiration in rabbits. These parameters were monitored using the wired telemetry system (Figure 5A & 5B). Healthy rabbits displayed a heart rate ranging from 190-200 beats per minute while rabbits with ALF showed a reduction in heart rate (140-150 beats per minute) (Figure 5C). Respiration rate in healthy rabbits was 75-105 breaths per minute while ALF rabbits showed a reduction (35 breaths per minute) (Figure 5D). The basal body temperature in healthy rabbits was 38-39°C, whereas rabbits with ALF had mild hypothermia with 37-38°C (Figure 5E). ECG was acquired using a single lead system (1^st^ and 2^nd^ panel, Figure 5F) and was superimposed for comparison. We show that animal with ALF had a prolonged Q-T interval and a lower QRS amplitude in comparison to healthy individual (Figure 5F, bottom panel). Waves Q through T measure the electrical activity in heart ventricle. Longer Q-T intervals indicate that it takes longer for the heart to contract and refill with blood before it beats again. Low QRS voltage indicates dampening of the signal due to excess fluid build-up in heart (pericardial/pleural effusion) or cardiomyopathy. This correlates to human clinical studies where similar changes in ECG have been observed in liver failure ^31^.

**Figure 5:**
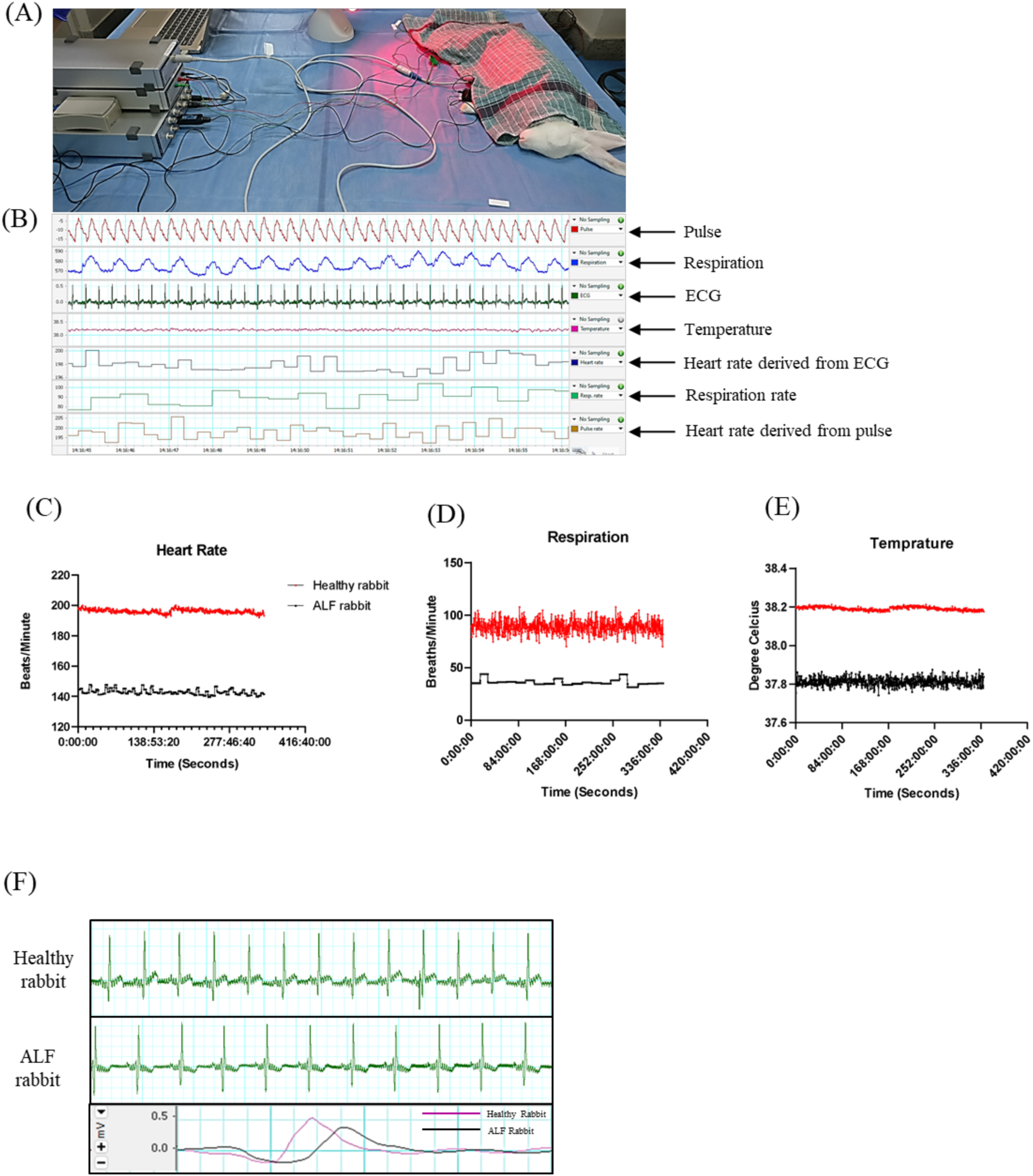
Effect of D-Galactosamine on vital signs in rabbits. (A) Representation of the anesthetized (35 mg/kg Ketamine and 5 mg/kg Xylazine) animal connected to the wired telemetry system (Power Lab C, AD Instruments) to capture the vital parameters. (B) Representation of the vitals acquired using Lab Chart 8 software. (C), (D), (E) and (F) show graphical representation of the vitals like Heart Rate, Respiration Rate, Temperature and ECG in heathy animal and animal with ALF (450mg/kg D-Galactosamine).

### NAC does not show any adverse effects in rabbits

As described earlier, NAC has been used as a hepatoprotective agent. Clinical & preclinical reports show a varying range of therapeutic dose of NAC being used ^32^. Eraslan G. et.al., have reported benefit from 500mg/kg NAC in aflatoxicosis in rabbits ^33^. There are minimal reports that have demonstrated the protective effects of NAC in non-acetaminophen drug induced ALF in rabbit model. The mechanism of action and/or the adverse effects of a high dose of NAC remains unclear. To understand this, the animal was injected with 500mg/kg NAC per day IM for 5 days during which blood sampling was performed every alternate day (Figure 6A). During this experiment, all the LFTs were found to be within the normal range. The levels of aminotransferases, bilirubin, serum albumin and clotting factors remained unaltered (Figure. 6B-6F). This is indicative of a healthy liver function thereby proving that NAC treatment does not cause any adverse effects on liver function. Overdose of NAC can lead to hemolysis and acute renal failure in humans ^34^. 500mg/kg NAC did not alter the levels of creatinine and urea, thus establishing safety of the administered dose (Figure 6G & 6H). The apparent behaviour of the animal remained normal during the experiment. All animals displayed normal feeding and drinking patterns and no fur loss was observed. In a nutshell, we hereby demonstrate that a dose of 500mg/kg NAC does not cause any adverse effects in rabbits.

**Figure 6:**
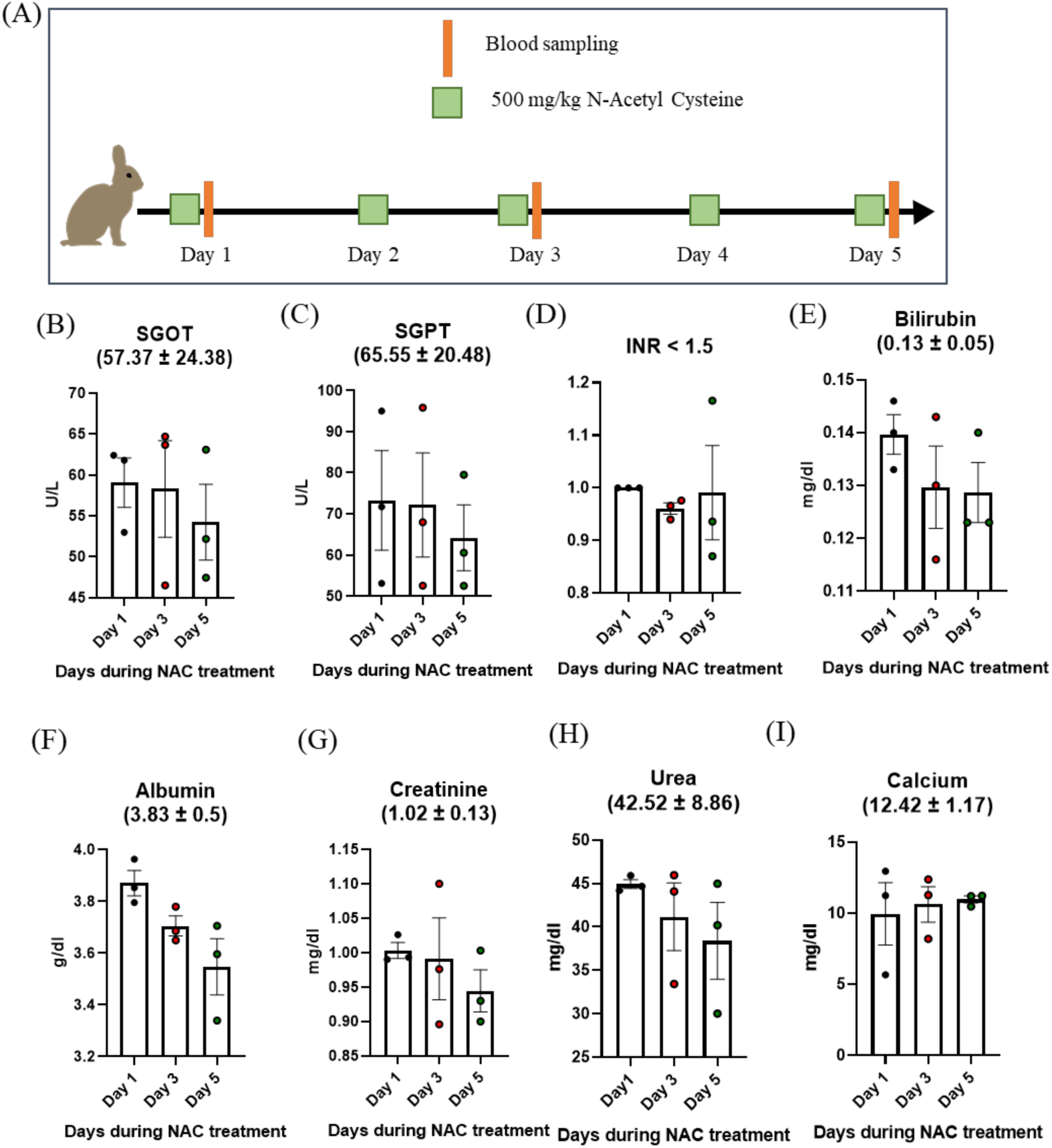
Evaluation of the safety of 500 mg/kg NAC in rabbits. (A) Schematic representation of the experimental strategy for determination of NAC toxicity *in-vivo*. (B), (C), (D), (E) and (F) Represent the liver function markers namely SGOT, SGPT, INR, Bilirubin and Albumin every 48 hours during NAC treatment (500 mg/kg every 24 hours for 5 days). (G) & (H) Represent the kidney function markers namely creatinine and urea every 48 hours during NAC treatment. (I) Represents the calcium level during NAC treatment. All the figures represent pooled data from N=3; n=3 wherein each point in graph represent data from one single animal.

### NAC protects against D-Galactosamine induced ALF and increases survival

Having established safety and the potency of NAC to suppress ROS *in-vitro*, we wanted to investigate the protective effects of NAC *in-vivo* in rabbits. The animals were treated with 500 mg/kg NAC every 24 hours for 5 days and on day 6 ALF was induced by IV administration of D-Gal in NAC and saline treated control groups (Figure 7A). We observed that NAC pre-treatment protects animals from drug induced ALF. The analysis of LFTs showed that control animals suffered from rapidly progressing liver damage that led to mortality. While the NAC pre-treatment protected the animals from severity and progression of ALF as indicated by remarkably lower ALT & AST levels (Figure 7B & 7C). Similarly, NAC pre-treated animals showed a lower INR of 1.6 compared to INR of 2.55 of controls (Figure 7D) and had normal albumin levels (Figure 7G). Most importantly, NAC pre-treatment provided a survival benefit i.e., the NAC treated cohort had a significantly higher survival of 77.8% compared to 33.3 % in controls (Figure 7E). Despite the significantly lower liver damage, the total bilirubin was high and comparable to controls (Figure 7F). This may be attributed to the longer half-life of bound bilirubin in blood (12-14 days). Reduced serum albumin production is often associated with hypocalcaemia ^35^. NAC treated animals showed only mild hypocalcaemia while the calcium levels in control group dropped markedly (Figure 7H). The blood urea levels were normal (Figure 7I), and all animals showed a mild increase in creatinine levels (Figure 7J). Overall, these results indicate that NAC pre-treatment can protect against severe liver injury *in-vivo* and increase survival.

**Figure 7:**
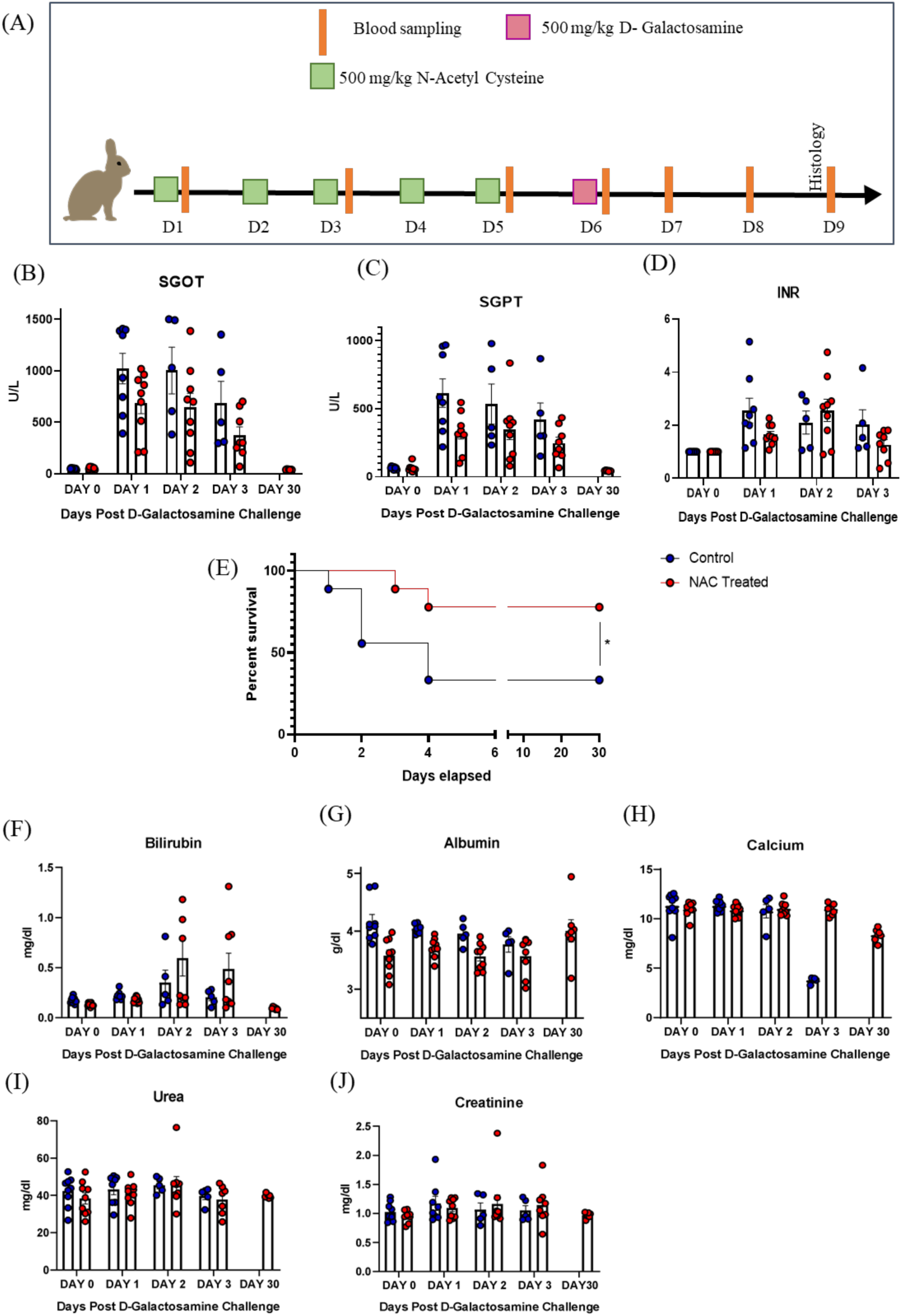
Protective effects of NAC against D-Galactosamine induced ALF in rabbits. (A) Schematic representation of the experimental strategy for illustrating the protective effects of NAC against ALF. (E) Represents the survival plot of animals treated with 500 mg/kg NAC and the vehicle control group after D-Gal administration. Statistics was performed using Log rank test where p*<0.05; N=3, n=3. (B), (C), (D), (F) and (G) Represents the liver function markers namely SGOT, SGPT, INR, Bilirubin and Albumin after D-Gal challenge in NAC treated and control groups. (H) Represents the calcium levels after D-Gal challenge in NAC and control groups. (I) & (J) Represent the kidney function markers namely creatinine and urea after D-Gal challenge in NAC treated and control groups. All the figures represent pooled data from N=3; n=3 wherein each point in graph represents data from one single animal.

### NAC pre-treatment protects the liver from D-Gal induced necrosis and inflammation

Having observed a clear survival benefit from ALF upon NAC treatment, a gross anatomical and histological examination was conducted. During necroscopy the liver from all control animals showed extensive damage (marked by discoloration). All animals from control cohort had necrotic lesions with haemorrhagic patches and hepatic portal inflammation. As per the ethical guidelines, the surviving animals from NAC pre-treatment group were sacrificed after 30 days. 4 out of 6 NAC treated animals displayed a completely normal liver architecture on Day 30 (Figure 8A, right panel), one animal was excluded from analysis due to secondary infection. There were considerably lesser necrotic lesions and lower inflammation compared to control cohort (Figure 8A & 8B). Table 3 and Table 4 shows the various histopathological parameters and their respective scores. All these results in total indicate the protective effects of NAC against ALF to the core.

**Figure 8:**
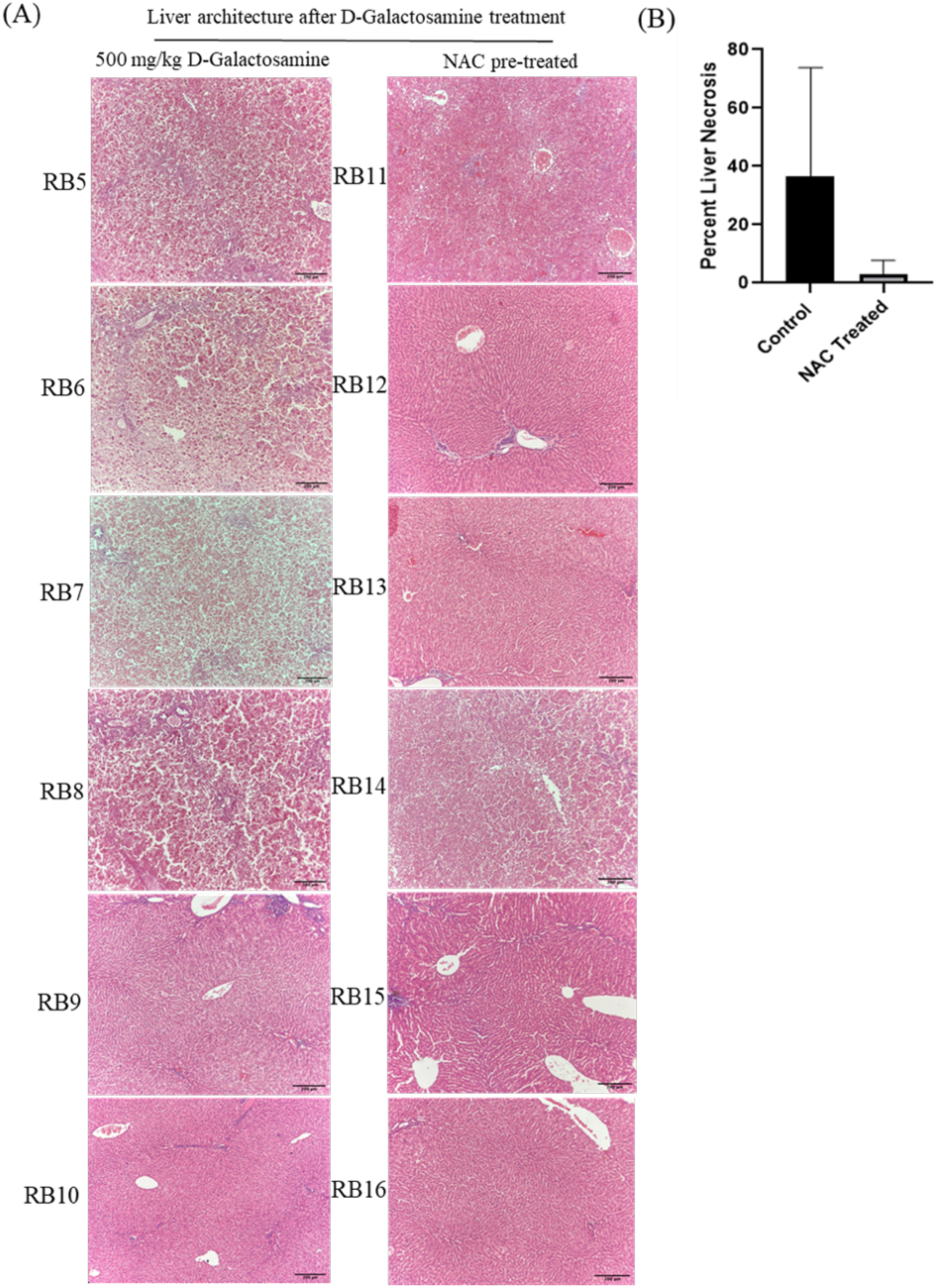
Histopathology evaluation of rabbit liver to establish the protective effects of NAC against D-Galactosamine induced ALF. (A) H&E staining representing the hepatocellular structure in control and 500 mg/kg NAC treated cohorts. Images were captured using a bright field microscope at 10X objective and analysed using Image J. Scale bar = 200 µm (B) Represents quantification of liver necrosis in control group and the surviving animals from NAC pre-treated cohort on Day 30 after D-Gal challenge.

**Table 4:** Histopathology Scoring of 500 mg/kg NAC pre-treated and 500 mg/kg D-Gal injected animals.

| **NAC-Treated Animals** |  |  |  |  |  |  |  |
| --- | --- | --- | --- | --- | --- | --- | --- |
| **Histology parameter** | **Location** | **Grade** |  |  |  |  |  |
|  |  | **RB11** | **RB12** | **RB13** | **RB14** | **RB15** | **RB16** |
| Steatosis |  | 5% | 1 % | 1% | 1% | 5% | 20% |
| Fibrosis | Periportal | 1 | 1 | 1 | 1 | 1 | 1 |
| Inflammation | Lobular | 1 | 1 | 1 | 1 | 1 | 1 |
| Inflammation | Portal | 0 | 0 | 0 | 0 | 0 | 0 |
| Necrosis |  | Absent | Absent | 5-8% | Absent | 5-10% | 15-20% |

## Discussion

ALF is a rare but lethal syndrome with no established treatments apart from OLT. The non-surgical treatments include the usage of antioxidant drugs like NAC which has been repurposed to treat liver failure in hospitals since its first use in APAP poisoning in 1977 ^36^. It has been firmly established to be safe and effective for this condition. Some randomized clinical trials showed that NAC increased transplant free survival and shorter hospital stay in NAI-ALF ^10,13.^ However, the evidence confirming the efficacy and safety of NAC with respect to NAI-ALF is lacking and the mechanism of action also remains unknown. Recent studies have shown that NAC can protect against genotoxic stress induced by cisplatin, tetrachlorobenzoquinone and hypoxia *in-vitro* ^37–39^. *In-vivo* studies report its protective action against carbon tetrachloride (CCl_4_), carbamazepine and diclofenac induced liver intoxication in rodent models ^40–42^. Hua Wang et al. demonstrated the hepatoprotective effects of NAC in endotoxin model of liver injury in D-Gal sensitized mice ^43^. Therefore, preclinical studies that demonstrate the protective effects of NAC are limited and have addressed this in rodent models only with respect to elevation of aminotransferases. The effects of NAC against D-Gal, a well-known hepatocyte toxin, remain unexplored till date. We performed a systematic study to explore the protective effects of NAC against D-Gal in hepatocyte cell line and *in-vivo* rabbit model (Figure 1). We established a dose dependent model of D-Gal induced toxicity in HepG2 cell line. NAC pre-treatment for 3 hours protected against D-Gal induced toxicity and improved cell survival. A dose as low as 100 µM NAC was sufficient to scavenge all the ROS, marking the saturation point for protection. Mortality in rodent models of ALF is less than 48 hours which does not provide a feasible timeframe for therapeutic interventions ^20,44^. The model demonstrated by us establishes 500 mg/kg D-Gal as the lethal dose that provides a time window of 48-96 hours. Blitzer et al. reported that the D-Galactosamine model of ALF in rabbits closely resembles human ALF in terms of plasma coagulation and EEG. In this model, we report the dose dependent kinetics of ALF progression in terms of all the LFTs as opposed to previous studies that have focused only the ALT and AST for evaluating liver damage. We observed that most of the LFTs peaked at 48 hours post intoxication. In addition, we observed that animals with ALF have lower heart rate, respiration and body temperature compared to healthy animals. ECG also showed longer Q-T interval and lower QRS amplitude like liver diseases in humans. These changes in animal vitals that are specific to ALF largely remain unknown and an in-depth characterization may serve as non-invasive yet crucial markers of disease progression ^45^. One of the key challenges with animal models of ALF is that they represent acute liver injury (ALI), which is less severe and therefore, does not mimic all the characteristics of ALF in humans ^46^. Despite this, it is crucial to test the therapeutic drugs in a pre-clinical setting. We demonstrate the protective effects of NAC against D-Gal induced liver injury *in-vivo.* Pre-treatment with safe doses of NAC for 5 days improved the survival of animals and effectively reduced severity of ALF within 3 days of intoxication. This reduction maybe due to ROS scavenging mechanism by replenishment of GSH pool *in-vivo* ^47^. NAC pre-treatment also improved all the LFTs upon D-gal intoxication compared to vehicle control. Interestingly, the total bilirubin levels remained elevated although the other LFTs showed a considerable reduction. In conclusion, our study reveals NAC as a potent hepatoprotective agent with no adverse effects at dose regimen of 500 mg/kg/day for 5 days. We show these hepatoprotective effects in unique D-Gal induced liver toxicity model in hepatocytes and New Zealand white rabbits. However, the safety and efficacy for long term use remains elusive and should be determined in future studies.

## Animal Ethics Statement

This study was approved and conducted as per the ethical guidelines by the Institutional Animal Ethics Committee (IAEC) with a registration number of 1803/PO/RcBi/S/2015/CPCSEA and project approval number of RR/IAEC/100-2022 & RR/IAEC/101-2022.

## Author Contributions

AD conceived the study. AD, TV, NK and AK performed experiments, analyzed data and prepared the manuscript. JG and MG provided critical input.

## Acknowledgements

The authors acknowledge the funding individuals and companies. Prof. Dipshikha Chakravortty’s laboratory is acknowledged for the equipment usage.

